# Where’s my mom? Resilient maternal preference in post-weaning male and female mice within a multi-chamber social behavior task

**DOI:** 10.1101/2025.03.05.641280

**Authors:** Maggie M. Slamin, Indra R. Bishnoi, Izabella M. Bankowski, Haley A. Norris, Evan A. Bordt

## Abstract

One of the earliest and most critical social bonds for many mammals is formed with their mother, who provides essential benefits for offspring development and survival. Growing evidence suggests that this social bond is retained even when animals gain independence, such as during the juvenile period immediately post-weaning. Here, we investigated whether juvenile (postnatal day (P)26) mice retain the ability to recognize and prefer their mothers post-weaning. We further investigated the strength of this bond using an acute immune activator. On P26, male and female C57BL/6J mice were intraperitoneally injected with the endotoxin lipopolysaccharide (LPS) or saline control (0.5 mg/kg). Four hours later, mice were subject to a five-chamber social preference task (the AGORA) containing their biological mother, a sex-matched novel mouse, a sex-matched sibling, a novel object, and an empty chamber. Our findings reveal that juvenile mice exhibit a strong maternal preference, significantly greater than chance and higher compared to any other social or non-social stimuli. While LPS exposure reduced the time spent investigating all stimuli, juvenile maternal preference was not significantly altered by LPS exposure. These effects were especially pronounced in females, while subtle shifts towards novel exploration began to emerge in males by P26. These results suggest that juvenile mice have a robust social preference for their mother that is resilient to early-life immune activation. Moreover, the novel multi-chamber task employed in the present study offered a more nuanced understanding of how social bonds evolve and vary across sex.

## 1. Introduction

One of the first social bonds most animals form is with their mother. This relationship carries several benefits. For example, maternal separation in pre-weaning mice (postnatal day [P]21-25) has been implicated in altered cognition and emotionality, including impairments in memory, increased anxiety-like and depressive-like behaviors, as well as increased exploratory-defensive behaviors (Nishi, 2020; Wang et al., 2020). Thus, a recognition of and preference for mothers early on is extremely beneficial for development and ultimately survival, which likely contributes to this preference being evolutionarily conserved in many mammals (Broad et al., 2006). Accordingly, mice have been found to recognize and prefer their mothers before weaning (Laham et al., 2021; Shimizu et al., 2022). After weaning, specifically during adolescence, a preference for novelty has been found to trump a maternal preference, even though mice recognize their mothers at this stage (Laham et al., 2021). However, the ability to recognize and prefer their mothers does not seem to disappear immediately after weaning in mice. During the juvenile period (post-weaning but prior to the pubescent stage of adolescence, ∼P25 to P30), mice continue to recognize and prefer their mothers in the three-chamber and two-choice social behavior tasks (Mogi et al., 2017; Stroobants et al., 2020). Consequently, juvenile mouse social behaviors may be more complex than previously understood.

Similar to other mammals, the juvenile period may be a transitionary stage in which mice shift from a reliance on maternal care to increased independence (Kelley et al., 2004; Spear, 2000). Thus far, studies using traditional social behavior assays like the three-chamber task (Mogi et al., 2017; Stroobants et al., 2020) have pointed towards a maternal preference in early juveniles.

However, these binary approaches cannot fully capture why juveniles may prefer their mothers. For example, in a three-chamber task, do mice display a maternal preference or an avoidance of a novel or unfamiliar stimulus? It is plausible that juvenile mice, still developing their defensive and risk-taking capacities, may opt for the safety of any familiar stimulus rather than risk interacting with an unfamiliar conspecific (Spear, 2000). This gap highlights the need for more nuanced paradigms to deepen our understanding of how juvenile mice navigate complex social relationships. Thus, in this study we employed a multi-social preference task to measure whether or not juvenile male and female mice display a maternal preference in the presence of other familiar and unfamiliar stimuli.

Furthermore, we explored the strength of juvenile social preference under aberrant conditions using lipopolysaccharide (LPS). The endotoxin LPS is a cell wall component of Gram-negative bacteria (King et al., 2009). LPS is a well-established tool to induce an acute inflammatory response, which results in sickness behaviors like altered social interaction (Dantzer & Kelley, 2007; Hart, 1988; Kavaliers et al., 2022). Immune activation induced via LPS is typically associated with social withdrawal, especially from an unfamiliar conspecific. However, emerging evidence indicates that some rodents, such as female rodents, exhibit social approach following infection, favoring a familiar social partner over a novel social stimulus (Kavaliers et al., 2022;

Muscatell & Inagaki, 2021). Presently, the effects of LPS on juvenile social behaviors are not well understood. Early-life or childhood immune activation has been linked to several neuropsychiatric disorders, such as schizophrenia, autism spectrum disorder, and major depressive disorder (Hoeijmakers et al., 2014; Ma et al., 2024; Mondelli & Vernon, 2019), which share a common dimension of aberrant social behaviors. Thus, characterizing early-life social behaviors, and potential disruptions to these behaviors, can provide significant insights into (a)typical social maturation.

In the present study, we examined juvenile social preference in a multi-chamber social preference task. We employed a five-chamber task, also known as the AGORA task, as it allows for the simultaneous measurement of social preference between five stimuli (Krueger-Burg et al., 2016). Further, we used this more sensitive assay to characterize the effects of LPS on these social behaviors during the juvenile period in male and female mice. Our findings reveal that juvenile mice exhibit a strong preference for their mother, significantly greater than chance and higher compared to other stimuli, including a novel mouse, a familiar sibling, a novel object, or an empty chamber. While LPS significantly reduced the time spent investigating each stimulus, the juvenile maternal preference was not significantly altered by LPS exposure, especially in female mice. These results suggest that juvenile mice have a robust social preference for their mother that is resilient to early-life immune activation.

## 2. Methods

### 2.1. Animals

Eight-week-old female wild-type C57BL/6J mice were purchased from Jackson Laboratories (Bar Harbor, ME: Stock #000664) (∼19.6 ± 1.2 grams upon arrival). Six harem breedings were conducted for two weeks in standard polypropylene cages (27 cm x 16 cm x 15.50 cm). Males were removed on the 14th day of breeding and dams were individually placed into new standard cages. All offspring were ear tagged for identification on P18. On P24, male and female offspring were weaned and group housed in standard cages with same-sex littermates. From P24, mothers were housed separately in their residing cages. All cages contained ad libitum access to water and food pellets (Prolab Isopro RMH 3000 Irradiated Pellets) in addition to Carefresh bedding and an igloo for environmental enrichment. All mice were housed in a colony room maintained at 23 ± 2°C with 50% humidity under a 12:12 light/dark cycle. All procedures were carried out during the light phase of the light/dark cycle (09:00–15:00 hr). All experiments were performed in accordance with the NIH *Guide to the Care* and *Use of Laboratory Animals* and approved by the Massachusetts General Hospital Institutional Animal Care and Use Committee.

### 2.2. Materials

#### 2.2.1. Lipopolysaccharide or saline injections

Lipopolysaccharide (from Escherichia coli serotype 0111: B4, Sigma-Aldrich, St. Louis, MO, USA) was dissolved in 0.9% sterile saline at 0.1 mg/ml. Mice were intraperitoneally administered either LPS or 0.9% saline control at a dose of 0.5 mg/kg 4 h prior to behavioral testing on P26. The selected route of administration, dose, and timing have been shown to be sufficient to induce sickness behaviors and changes in social behavior in male and female mice (Aries et al., 2023; Corona et al., 2010; Hines et al., 2013; Ohgi et al., 2013; Vichaya et al., 2020). Experimental animals were randomly assigned to either the LPS or saline control groups. However, mice within the same cage received the same injection. To avoid litter effects, only two mice of each sex were randomly assigned from each litter to be experimental animals. Thus, the current study consisted of four groups: female-saline (n = 15), female-LPS (n = 13), male-saline (n = 12), male-LPS (n = 12).

#### 2.2.2. Behavioral apparatus

The AGORA social preference task (Ugo Basile Catalog #46573) is a five-chambered apparatus (61 cm x 58 cm x 25 cm). Within the chamber, there is a central arena (50 cm x 50 cm). Attached to the central arena are 5 removable chambers (each 13 cm x 11.5 cm). The central arena is separated from the 5 chambers through removable plexiglass dividers which contain 41 holes (each with a diameter of 0.8 cm) at the bottom of every divider to allow animals to identify odors during social exploration (see Krueger-Burg et al., 2016).

### 2.3. Experimental procedures

On P25, offspring and mothers were habituated in the behavioral testing room for 1 h with dimmed lights. Cages were covered with a white tarp to minimize the ability to see the behavioral testing apparatus. Following habituation to the room, experimental mice were habituated to the arena of the apparatus for 7 mins. Novel mice, siblings, and mothers were habituated to a randomly assigned chamber for 7 mins. Multiple novel mice, siblings, and mothers were allowed to habituate within the same round of habituation in separate chambers. The apparatus was cleaned with Peroxigard and allowed to completely dry between each round of habituation. On P26, experimental male and female offspring received an intraperitoneal injection of saline or LPS. Following injection, they were placed back into their home cages and returned to their colony room. Mice were moved to the behavioral testing room 1 h prior to behavioral testing to habituate them to the room. After this period, all stimuli were placed into the chambers of the AGORA (novel stimulus, sibling, mother, novel object, and an empty chamber). Locations of the stimuli were randomly assigned and varied for each trial. Exactly 4 h post injection, an experimental animal was placed into the center of the arena of the AGORA task and was allowed to explore the arena for 7 mins. The apparatus was cleaned with Peroxigard and allowed to completely dry between each trial. All trials were video recorded. The following behavioral outcomes were manually scored by a blinded observer using Solomon Coder: time spent investigating the novel mouse, time spent investigating the familiar cage mate sibling, time spent investigating the biological mother, time spent investigating the novel object, time spent investigating the empty chamber, and non-investigation time. Distance moved and velocity were obtained using Ethovision 17.5 (Noldus).

### 2.4. Statistical analyses

A two-way analysis of variance (ANOVA) was used to analyze percentage body weight change, distance moved, velocity, and non-social interaction time, with Treatment (at two levels: LPS or saline) and Sex (at two levels: male or female) as the independent variables. Significant main effects and interactions were further examined using Tukey’s honestly significant difference (HSD) post hoc test. A repeated measures ANOVA was used to analyze the time spent investigating each chamber during the multi-chamber social preference task. The within-subject factor was Chamber (at five levels: maternal, novel, sibling, object, and empty), and the between-subject factors were Treatment (LPS or saline) and Sex (male or female). Greenhouse–

Geisser corrections were applied where Mauchly’s test of sphericity was violated. Significant effects were further examined using the Bonferroni post hoc test. A one-sample t-test was used to compare the percentage of time spent investigating each chamber to the chance criterion of 20%. All data passed the Shapiro-Wilk Test for normality. A two-way ANOVA was conducted to analyze the percentage of time spent investigating each chamber, with Treatment (LPS or saline) and Sex (male or female) as independent variables. Significant effects were further examined using Tukey’s HSD post hoc test. All statistical tests employed a significance criterion of α = 0.05. All data are reported as mean ± SEM. All statistics and data visualization were performed using GraphPad Prism Version 10. Individual points on graphs represent individual experimental mice.

## 3. Results

### 3.1. 0.5 mg/kg LPS induced sickness behaviors in male and female juvenile mice

To assess whether or not 0.5 mg/kg of LPS altered behaviors in P26 male and female mice, we measured standard sickness responses following treatment with LPS (Dantzer & Kelley, 2007; Muscatell & Inagaki, 2021) compared to saline treated controls.

A sickness response to LPS administration is a decrease in body weight. Using a two-way ANOVA, we found a significant main effect of treatment on % body weight change (*F*(1, 48) = 38.580, *p* < .0001) (Figure 1A). Mice treated with LPS demonstrated a significant decrease in body weight 4 h following treatment compared to saline-treated mice. A significant interaction effect was also found (*F*(1, 48) = 38.580, *p* < .01). Male mice treated with LPS weighed significantly less than saline-treated male mice (*p* < .0001). Furthermore, saline-treated female mice had a significantly lower % body weight change than males treated with saline (*p* < .05). Male and female mice treated with LPS did not significantly differ in their % body weight change, suggesting that LPS impacted male and female mice similarly. We did not find a significant main effect of sex.

**Figure 1.**
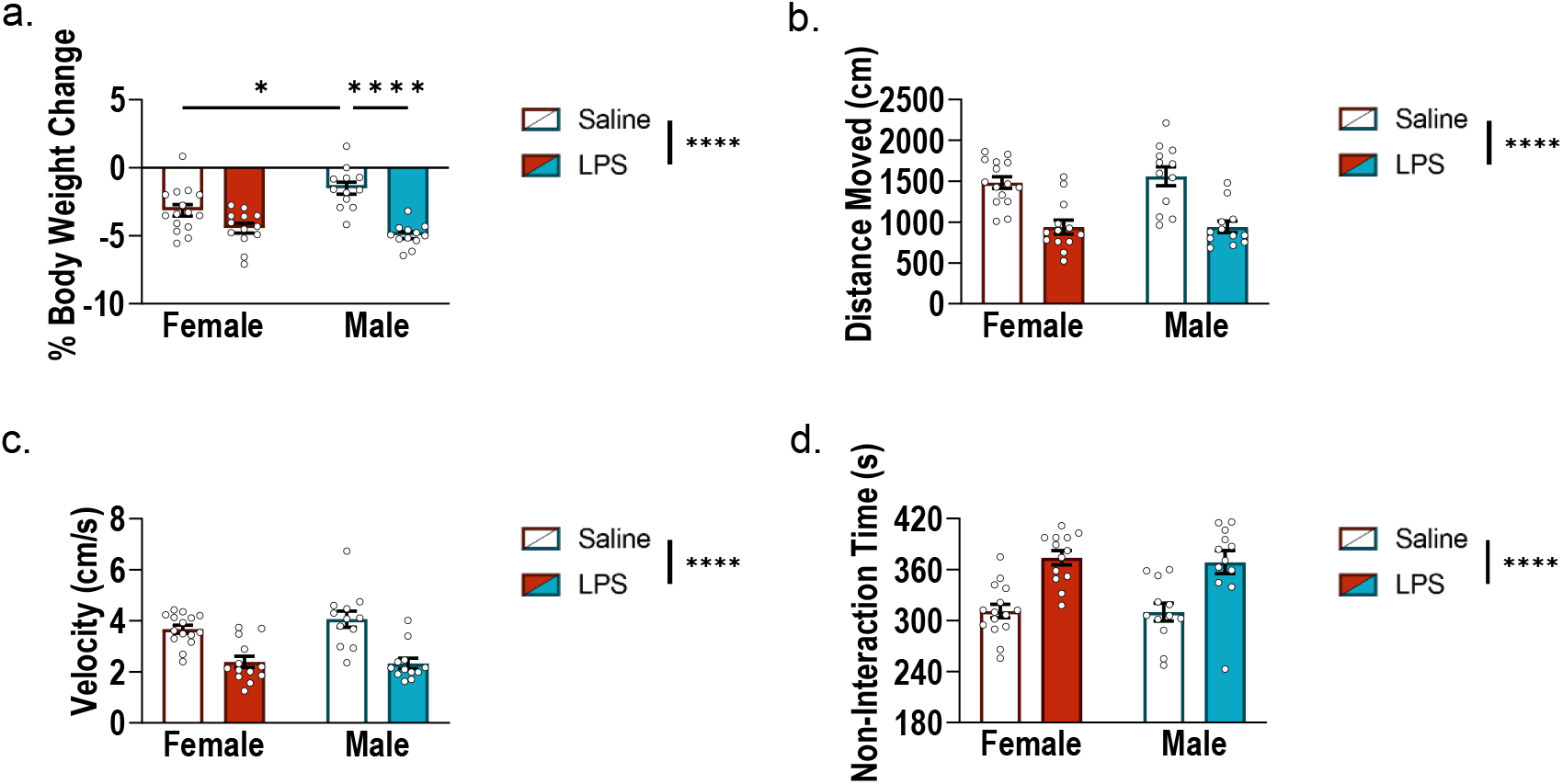
0.5 mg/kg LPS induces sickness behaviors in male and female juvenile mice. **(a)** Percent body weight change 4 hours post-treatment in P26 male and female mice administered LPS or saline. LPS-treated mice exhibited a significant reduction in body weight compared to saline-treated controls (*p* < .0001). Male mice treated with LPS weighed significantly less than saline-treated males (*p* < .0001). Saline-treated females had a significantly lower % body weight change than saline-treated males (*p* < .05). **(b)** Total distance moved (cm) and **(c)** movement velocity (cm/s) in the AGORA task. LPS exposure significantly decreased both the distance traveled and the velocity of movement compared to saline controls (*ps* < .0001). **(d)** Non-interaction time (s) in the AGORA task. LPS-treated mice spent significantly more time not interacting with any chamber compared to saline-treated mice (*p* < .0001). \**p* < .05, \*\*\*\**p* < .0001. Data are presented as mean ± SEM.

The administration of LPS also resulted in other sickness responses including reduced locomotion and increased isolation or non-interaction time. A two-way ANOVA revealed a significant main effect of treatment on distance moved (*F*(1, 48) = 45.640, *p* < .0001) (Figure 1B) in addition to velocity of movement (*F*(1, 48) = 43.910, *p* < .0001) (Figure 1C). Compared to saline, LPS significantly reduced both the distance and velocity moved by mice compared to saline controls. Finally, a significant main effect of treatment was observed on non-interaction time (*F*(1, 48) = 36.25, *p* < .0001) (Figure 1D), as the administration of LPS significantly increased time spent in isolation (not investigating any chamber in the AGORA) compared to saline-treated mice. No significant main effect of sex or interaction effects were found.

### 3.2. LPS reduced the duration of social investigation

To investigate the duration of social and non-social investigation in juvenile mice, we exposed P26 mice to a multi-chamber social preference task wherein they were allowed to explore a five-chamber task for 7 mins with the opportunity to investigate chambers containing their mother, a novel age- and sex-match mouse, their sibling, a novel object, or an empty chamber. We assessed the duration that male and female mice who were administered either saline or LPS investigated each of these stimuli (Figures 2, 3). A repeated measures ANOVA revealed a significant main effect of treatment on the time spent investigating each chamber (*F*(1, 48) = 36.246, *p* < .0001). Compared to saline, the administration of LPS significantly decreased the time spent investigating each chamber (Figure 2). A significant main effect of chamber was also found (*F*(2.011, 96.54) = 23.561, *p* < .0001). Mice spent significantly more time investigating the maternal chamber compared to all other chambers (*ps* < .0001). Mice also spent significantly more time investigating the novel chamber compared to the object and empty chambers (*ps* <.05), but not the maternal or sibling chambers. Finally, mice spent more time investigating the sibling chamber compared to the empty chamber (*p* < .05) (Figure 3).

**Figure 2.**
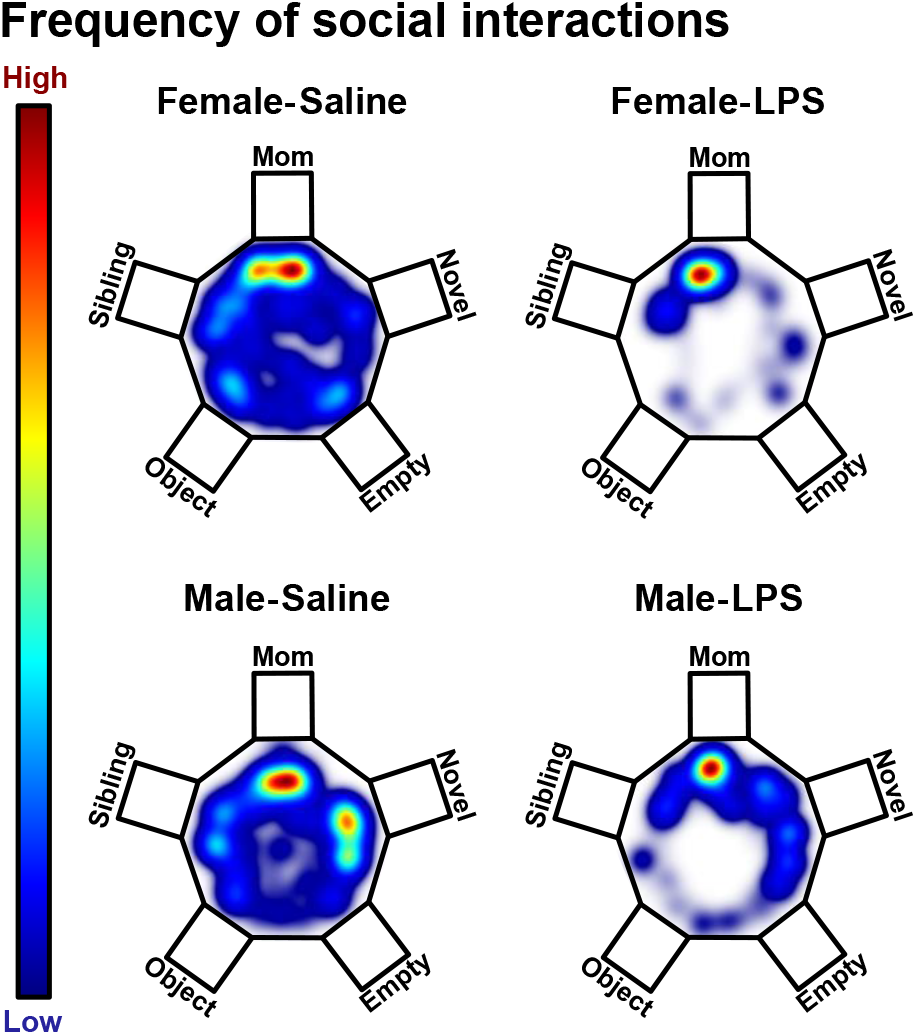
Representative heatmaps illustrating the frequency of social interactions in juvenile mice. Heatmaps depict spatial distribution of interactions across groups (Female-Saline, Female-LPS, Male-Saline, and Male-LPS) within the multi-chamber social preference AGORA task. The chambers are labeled to represent maternal, novel, empty, object, and sibling stimuli across groups; however, the placement of these stimuli was randomized across trials to prevent spatial biases. Warmer colors (red) indicate areas of higher interaction, while cooler colors (blue) represent areas of lower interaction.

**Figure 3.**
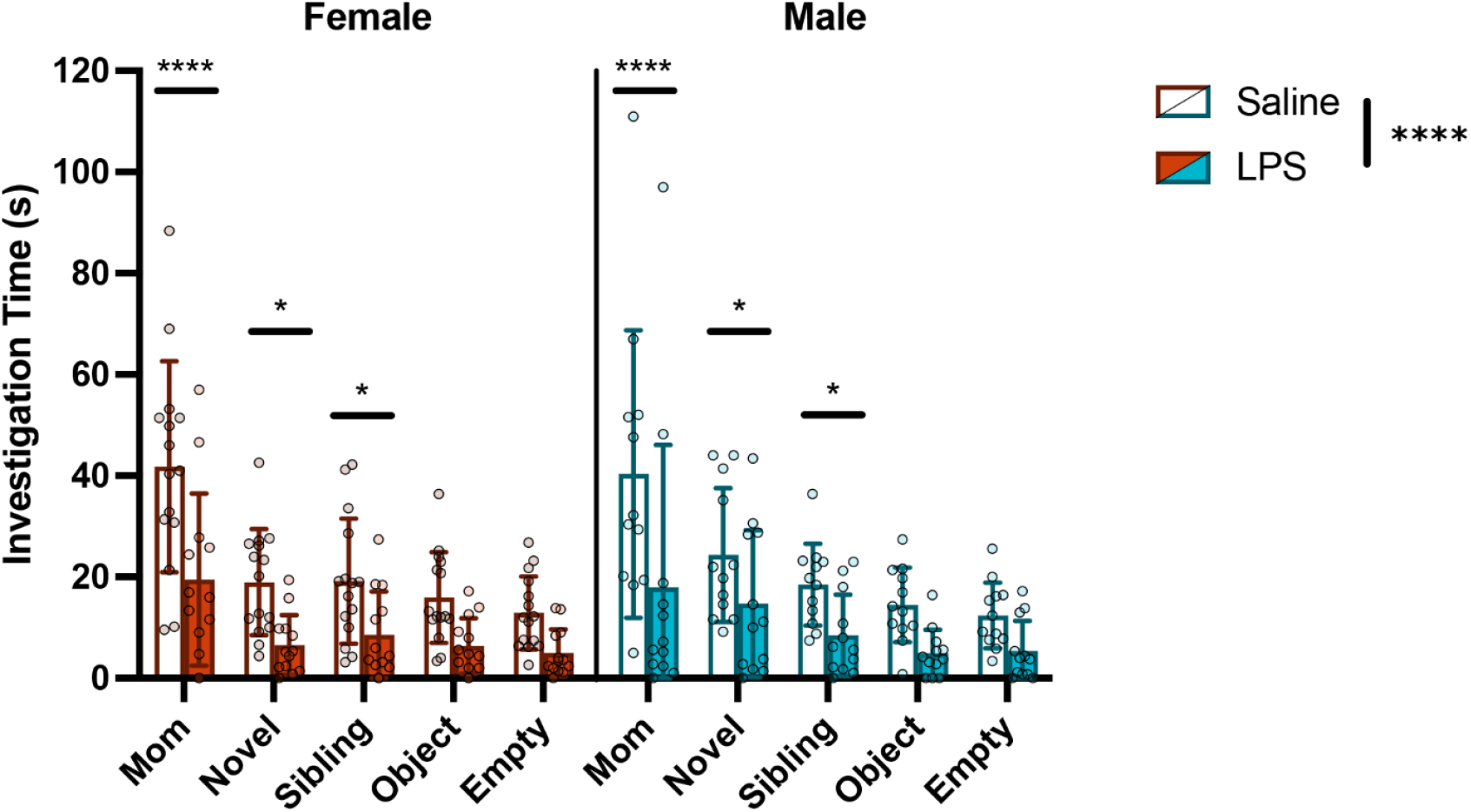
LPS reduces investigation of social and non-social stimuli in juvenile mice. Male and female P26 mice administered LPS or saline were tested in a multi-chamber social preference task for 7 mins. Mice had the opportunity to investigate chambers containing their mother, a novel age- and sex-matched mouse, their sex-matched sibling, a novel object, or an empty chamber. LPS-treated mice spent significantly less time investigating all chambers compared to saline-treated controls (*p* < .0001). Mice spent significantly more time investigating the maternal chamber compared to all other chambers (*p* < .0001). Mice also spent significantly more time investigating the novel mouse compared to the object and empty chambers (*p* < .05). Investigation of the cage mate sibling chamber was significantly greater than the empty chamber (*p* < .05). \**p* < .05, \*\*\*\**p* < .0001. Data are presented as mean ± SEM.

### 3.3. Juvenile mice spend a greater percentage of their time engaging in maternal investigation compared to chance

Next, we aimed to understand juvenile social preference in the AGORA by comparing the percentage of time spent investigating each chamber to the chance criterion of 20% (indicated using a horizontal grey line, see Figure 4). As such, a one-sample t-test was used to evaluate whether the mean of percentage of time spent investigating each chamber differed from chance. One-sample t-test revealed that saline-treated female mice (*M* = 36.75, *SEM* = 3.283, *t*(14) = 5.101, *p* < .001), LPS-treated female mice (*M* = 38.04, *SEM* = 6.506, *t*(12) = 2.772, *p* < .05), and saline-treated male mice (*M* = 33.81, *SEM* = 4.249, *t*(11) = 3.250, *p* < .01) spent a greater percentage of time investigating the maternal chamber compared to chance. However, LPS-treated male mice did not spend a greater percentage of time investigating the maternal chamber. Juvenile mice did not differ in the percentage of time spent investigating the novel or sibling chambers compared to chance. However, LPS-treated female mice (*M* = 13.35, *SEM* = 2.932, *t*(12) = 2.269, *p* < .05) and saline-treated male mice (*M* = 14.46, *SEM* = 2.129, *t*(11) = 2.603, *p* < .05) spent a lower percentage of time investigating the object chamber. The saline-treated female mice and LPS-treated male mice did not statistically differ in their percentage of time spent investigating the object chamber compared to chance. Finally, both female (*M* = 11.44, *SEM* = 1.212, *t*(14) = 7.066, *p* < .0001) and male (*M* = 11.35, *SEM* = 1.449, *t*(11) = 5.972, *p* < .0001) saline-treated mice spent a lower percentage of time investigating the empty chamber compared to chance. LPS-treated female and male mice did not statistically differ in their percentage of time spent investigating the empty chamber compared to chance (Figure 4).

**Figure 4.**
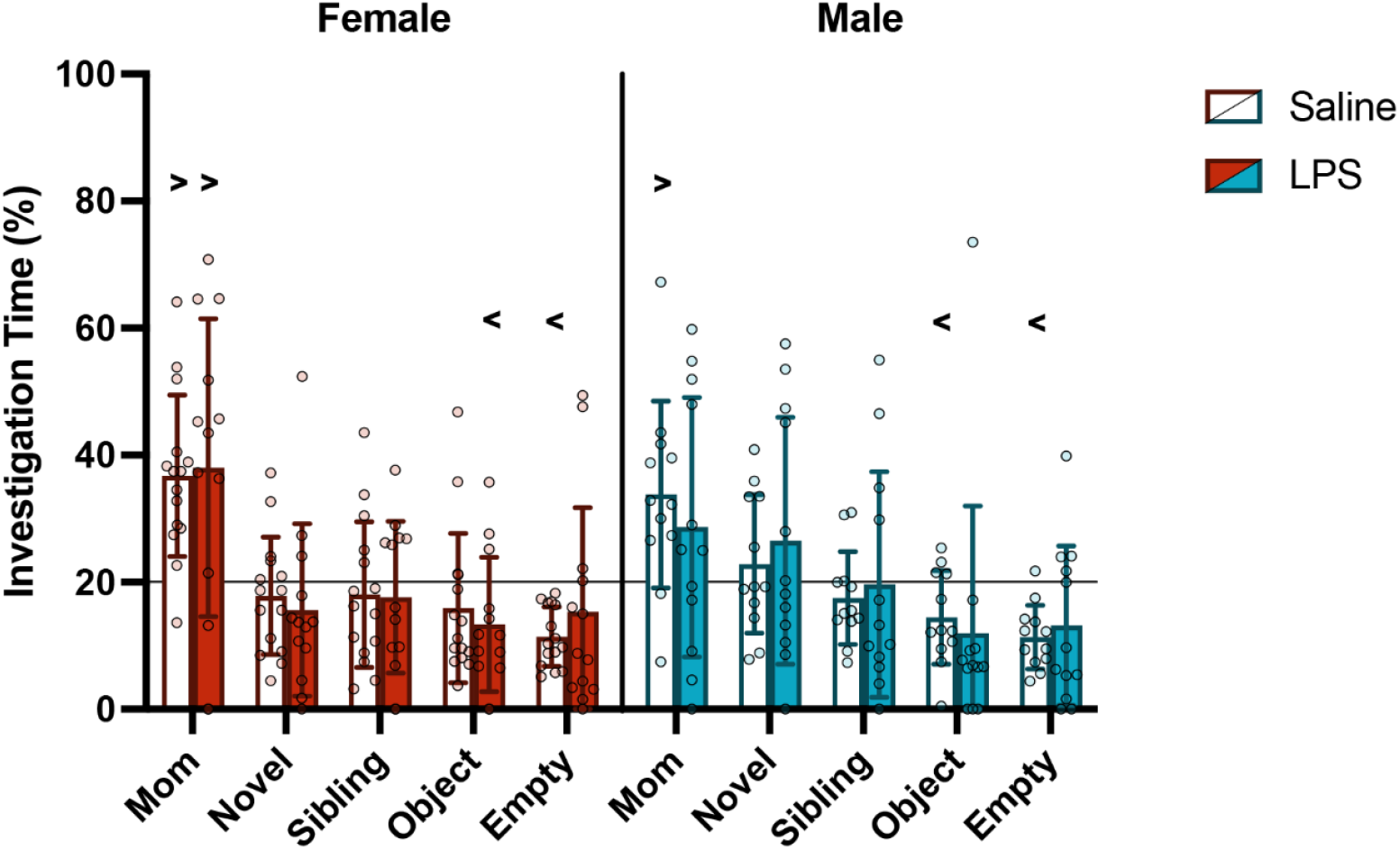
Juvenile mice preferentially investigate their mother compared to chance. The percentage of time spent investigating each chamber was calculated as a proportion of total investigation time. Percentage of time spent investigating each chamber in the AGORA social preference task was compared to a 20% chance criterion (indicated by a horizontal gray line). Female-saline mice, female-LPS mice, and male-saline mice spent a significantly greater percentage of time investigating the maternal chamber compared to chance. LPS-treated female mice and saline-treated male mice spent significantly less time investigating the object chamber compared to chance. Finally, both saline-treated female and male mice spent significantly less time investigating the empty chamber compared to chance. ^>^significantly greater than chance (*p* < .05) ^<^significantly less than chance (*p* < .05). Data are presented as mean ± SEM.

### 3.4. Juvenile males demonstrate a greater preference for novel stimuli compared to juvenile females

Consistent with previous studies, (Dantzer & Kelley, 2007; Muscatell & Inagaki, 2021), LPS treatment decreased locomotion and total investigation time (Figures 1b-c, 3). Therefore, to determine whether LPS impacted preference for which chamber to investigate, regardless of total investigation time, we evaluated the percentage of time spent investigating each chamber (Figure 5). The percentage of time spent investigating each chamber was calculated by dividing the percentage of time spent investigating each chamber by all chambers (e.g., % time investigating maternal chamber /all investigation) and analyzed using two-way ANOVAs.

**Figure 5.**
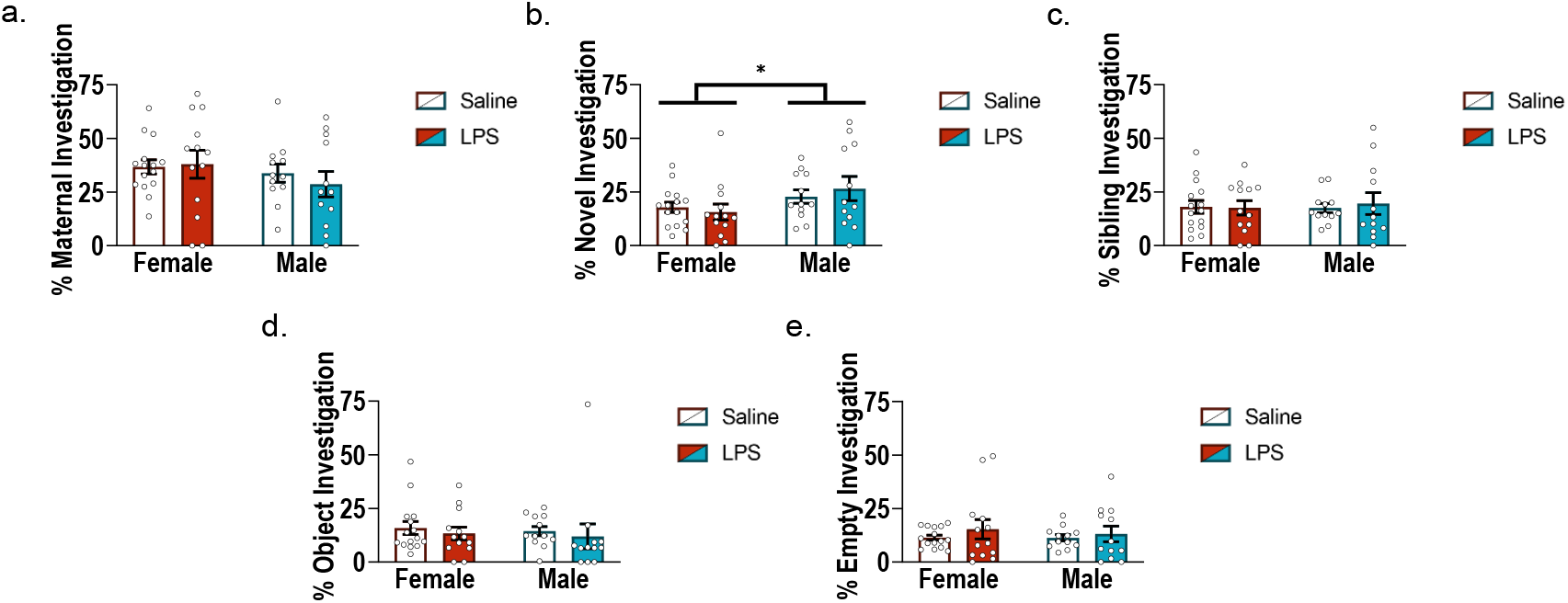
Juvenile males demonstrate a greater preference for novel social stimuli compared to females. The percentage of time spent investigating **(a)** the maternal **(b)** novel sex- and age- matched mouse **(c)** sex-matched sibling **(d)** novel object or **(e)** empty chamber was calculated as a proportion of total investigation time. **(b)** Male mice spend a significantly greater percentage of time investigating the novel stimulus mouse compared to female mice (*p* < .05). \**p* < .05. Data are presented as mean ± SEM.

Statistical analyses revealed no significant main effect of sex, treatment, or interaction effect on the percentage of time spent investigating their biological mother (Figure 5a). A significant main effect of sex was found for the novel chamber (*F*(1, 48) = 4.436, *p* < .05). Compared to the female mice, the male mice spent a significantly higher percentage of time investigating the novel stimulus mouse (Figure 5b). No significant main effect of treatment or interaction effect were found for the novel chamber. No significant main effect of sex, treatment, or interaction effect were found in the percentage of time spent investigating a sex- and age-matched sibling, an object, or the empty chamber (Figure 5c-e).

## 4. Discussion

Our findings demonstrate that juvenile mice recognize and exhibit a pronounced maternal preference, which surpasses their preference for any other tested stimuli including novel mice, siblings, novel objects, or empty chambers. This maternal preference persists despite LPS-induced immune activation and sickness behaviors (see Figure 2). Notably, while male mice showed a diminished preference for their mother following LPS treatment compared to chance, LPS-treated female mice continued to spend a greater percentage of time investigating their mothers, illustrating an especially resilient maternal preference in females. These findings suggest that juvenile mice, especially females, have a robust social preference for their mother at P26 that is resilient to the effects of early-life LPS-induced infection.

Both female and male mice displayed sickness responses following LPS treatment, such as altered social behaviors, reduced locomotion, and a reduction in body weight (Dantzer & Kelley, 2007; Muscatell & Inagaki, 2021). Stronger sickness responses were found in male mice, including greater time spent in isolation, reduced locomotion, and decreased body weight (see Figures 1, 2). These findings suggest that male mice may display a more pronounced shift in behavior in the AGORA 4 hours following LPS treatment. These findings are in line with findings in other social behavior tasks and tests of locomotion, wherein females display less evident sickness responses after LPS treatment (Bishnoi et al., 2021; Tenk et al., 2008; Yamamoto et al., 2025), though the effects on social behavior may be more complex (Decker Ramirez et al., 2023; Smith et al., 2020). Additionally, female rodents typically display greater stress-related responses at baseline (Goel et al., 2014), thus potentially depressing the magnitude of the difference in body weight between female saline and LPS groups, which can be altered due to the stress response rather than solely immune activation in rodents (Marin et al., 2007).

LPS-treated mice displayed a significant withdrawal from all stimuli, both social and non-social. However, based on the observed sickness responses, what appears to be social withdrawal from all stimuli may rather be a general reduction in locomotion. This is supported by our exploration of what LPS-treated juvenile mice do in the time they are investigating the chambers (i.e., the percentage of time spent investigating chambers). Compared to saline-treated mice, we found that LPS-treated female and male mice displayed a similar preference profile, with the strongest preference also being for their mothers. This finding suggests LPS-treated mice in the current study did not lose social recognition and preference per se. Instead, the decrease in exploratory behaviors could be due to a general suppression of locomotion that often accompanies the acute inflammatory response following LPS exposure (Bishnoi et al., 2022; Dantzer & Kelley, 2007).

An intriguing finding in the present study was that male mice spent more time investigating the novel sex- and age-matched mouse than the female mice, regardless of treatment. This finding suggests that, while male mice may recognize their mothers (as indicated by their overall greater exploration of the maternal chamber), they may begin to shift their preference towards novelty earlier than female conspecifics. Notably, this preference for the novel mouse did not differ from chance, indicating that their preference may begin shifting but is not completely shifted towards novelty at P26 when weaned on P24. This early inclination towards novelty in males is well-aligned with their developmental stage. The late juvenile/early adolescent stages are characterized by a period of heightened novelty seeking, which is crucial to facilitate the transition from dependence to independence (Kelley et al., 2004; Spear, 2000). Specifically for males, it may be adaptive to begin novelty seeking and transitioning towards independence earlier to reduce competition and inbreeding (Holekamp et al., 1984; Pusey, 1987). Indeed, sex differences in gaining independence are observed across mammalian species. In the wild, female mice remain in the neonatal nest longer than males (Schlegel & Barry, 1991). Further, other rodents, like Belding’s ground squirrels, display common mammalian sex-biased pattern in which males disperse with much greater probability, while females rarely leave their natal areas (Holekamp et al., 1984). This sex-dependent pattern has been found to be gonadal hormone-dependent. Female Belding’s ground squirrels treated with testosterone propionate within 36 h of birth dispersed away from their mothers close to male rates (Holekamp et al., 1984). Thus, our finding that 26-day-old male mice spent a greater percentage of time investigating the novel mouse compared to females aligns with the sex-specific developmental shifts which are beginning to occur at this developmental stage.

The ability of juvenile male and female mice to recognize and prefer their biological mothers aligns with previous studies conducted in two-choice and free-roaming social tasks showing that post-weaning mice retain maternal bonds (Laham et al., 2021; Mogi et al., 2017; Stroobants et al., 2020). Importantly, the five-chamber AGORA task allowed for the more nuanced understanding of these behaviors. For example, we were able to simultaneously control for cage-effects by including a familiar cage-mate sibling in this task. Our findings suggest that it was not the case that any littermate induced a social preference, but rather that social preference was targeted toward mothers specifically. Additionally, we were able to highlight the potential starting point for the transition towards a preference for novel social stimuli in male mice.

Finally, using the five-chamber task, we found that while LPS treatment significantly reduced social exploration and increased isolation, social preference (measured by the relative percentage of time spent investigating stimuli) in LPS-treated mice remained comparable to that of saline-treated mice.

Given the present findings, future research should map shifts in social preference (e.g., when novelty preference develops and subsides) in the context of developmental maturation. The multi-chamber AGORA task is particularly suited for these explorations as it allows for the simultaneous addressing of confounds (e.g., an avoidance of familiar over a novelty preference), which are difficult to discern in traditional social preference assays. Moreover, while researchers are now consistently seeing this robust maternal preference, the neurobiology underlying this preference is yet to be elucidated. Finally, the AGORA task will also aid in future research exploring the behaviors of the interacting partner, such as the preference of mothers towards or away from saline or LPS-treated pups. However, it is important to note that social dynamics vary significantly not just based on sex and age but also other factors like species and strain. Our work, focused on C57BL/6J mice, provides valuable insights into the developmental and behavioral dynamics of this particular strain. Other strains of mice, such as BALB/c mice, have been found to demonstrate lower sociability (Sankoorikal et al., 2006). In contrast, several strains of rats demonstrate greater sociability than mice (Hudgens et al., 1968; Kummer et al., 2014). Comparable studies are essential to determine the generalizability of the current findings.

This study demonstrates that post-weaning juvenile mice exhibit a strong and selective preference for their biological mothers over other social and non-social stimuli, a preference that remains intact even under immune challenge. These findings challenge the notion that mouse social behavior is guided by novelty-seeking and highlight the endurance of early maternal bonds during the juvenile period. The results also highlight the importance of considering sex differences in the developmental timing of social behaviors. By leveraging a multi-chamber behavioral paradigm, our study offers a more comprehensive perspective on the complexity of juvenile social decision-making than traditional binary assays. Future research should continue to explore the neurobiological underpinnings of these behaviors, which could offer insights into typical and atypical social development.

## Funding

This work was supported by R01 HD110467 to EAB as well as the Robert and Donna Landreth Family Foundation. IRB was supported by a Landreth Research Fellowship through the Robert and Donna Landreth Family Foundation.

## Notes

### Competing Interest Statement

The authors have declared no competing interest.

